# Genetically distant bacteriophages elicit unique genomic changes in *Enterococcus faecalis*

**DOI:** 10.1101/2021.10.28.466302

**Authors:** Cydney N. Johnson, Dennise Palacios Araya, Viviane Schink, Moutusee Islam, Mihnea R. Mangalea, Emily K. Decurtis, Tuong-Vi Cindy Ngo, Kelli L. Palmer, Breck A. Duerkop

**Affiliations:** Department of Immunology and Microbiology, University of Colorado School of Medicine, Aurora, CO, USA, 80045; Department of Biological Sciences, University of Texas at Dallas, Richardson, TX, USA, 75080

**Keywords:** Bacteriophages, *Enterococcus faecalis*, coevolution, comparative genomics

## Abstract

The human microbiota harbors diverse bacterial and bacteriophage (phage) communities. Bacteria evolve to overcome phage infection, thereby driving phage evolution to counter bacterial resistance. Understanding how phages promote genetic alterations in medically relevant bacteria is important as phages continue to become established biologics for the treatment of multidrug-resistant (MDR) bacterial infections. Before phages are used as standalone or combination antibacterial therapies, we must obtain a deep understanding of the molecular mechanisms of phage infection and how host bacteria alter their genomes to become resistant. We performed coevolution experiments using a single *Enterococcus faecalis* strain and two distantly related phages, to determine how phage pressure impacts the evolution of the *E. faecalis* genome. Whole genome sequencing revealed mutations previously demonstrated to be essential for phage infection. We also identified mutations in several genes previously unreported to be associated with phage infection in *E. faecalis*. Intriguingly, there was only one shared mutation in the *E. faecalis* genome in response to each of the two phages tested, demonstrating that infection by genetically distinct phages results in different host responses. This study shows that infection of the same host by disparate phages leads to evolutionary trajectories that result in distinct genetic changes. This implies that bacteria respond to phage pressure through host responses that are tailored to specific phages. This work serves as the basis for the study of *E. faecalis* genome evolution during phage infection and will inform the design of future therapeutics, such as phage cocktails, intended to target MDR *E. faecalis*.

**IMPORTANCE:** Studies characterizing the genome evolution of bacterial pathogens following phage selective pressure are lacking. Phage therapy is experiencing a rebirth in Western medicine. Such studies are critical for understanding how bacteria subvert phage infection and how phages evolve to counter such mutations. This study utilizes comparative genomic analyses to demonstrate how a pathogenic strain of *Enterococcus faecalis* responds to infection by two genetically distant phages. We show that genetic alterations in the *E. faecalis* genome accumulate in a manner that is specific to the infecting phage with little to no overlap in shared fixed mutations. This suggests that bacterial genome evolution in response to phage infection is uniquely tied to phage genotype, and sets a precedence for investigations into how phages drive bacterial genome evolution relevant to phage therapeutic applications.

## INTRODUCTION

*Enterococcus faecalis* is a Gram-positive bacterium naturally residing as a commensal in the gastrointestinal tracts of animals, including humans (1). Immune suppression and/or antibiotic treatment can cause *E. faecalis* to outgrow and become the dominant member of the microbiota, leading to life-threatening opportunistic infections (2). Strains of *E. faecalis* and *Enterococcus faecium* have acquired traits that allow them to survive host and environmental stresses, contributing to their success as pathogens (3, 4). The overuse of antibiotics in both medical and agricultural settings has played a large part in enterococcal pathogenesis by driving multidrug-resistant (MDR) phenotypes (5, 6). As MDR *E. faecalis* infections continue to persist worldwide, there is a need to find alternative therapeutics capable of bypassing existing modes of antibiotic resistance (7–9).

Bacterial viruses, bacteriophages (phages), exist in high numbers in the intestinal tract where they infect and sometimes kill host bacteria, likely influencing the structure of the microbiota (10–12). Due to their narrow host specificity and ability to lyse bacteria, phages are becoming an essential resource for the treatment of MDR bacterial infections (13). Phage therapy offers many advantages over traditional antibiotics. For example, specificity can be tailored to target only the desired bacteria, leaving native microbes largely unaffected (14, 15). Additionally, phage replication is restricted to the abundance of the host, thus upon host exhaustion phages are depleted from the population (16). In contrast, conventional antibiotics lack specificity, killing resident bacteria, and the compounds can remain in the patient after the infection has cleared (17). There is an ever-growing repertoire of phages that infect *E. faecalis* (18), making these promising candidates for phage therapy.

The development of successful phage therapies will require a complete understanding of the genetic interactions between phages and bacteria. Although phage therapy holds promise for the treatment of *E. faecalis* infections (19, 20), the molecular mechanisms of enterococcal phage infection and the bacterial host response to phage infection are understudied. Phage tail protein-receptor interactions underpin the molecular basis for phage strain specificity of the bacterial cell surface (21, 22). To date, only the transmembrane protein PIP_EF_ (phage infection protein of *E. faecalis*) has been identified as a bona fide enterococcal phage receptor (23). Both phages VPE25 and VFW bind to *E. faecalis* through cell surface polysaccharides, and infection proceeds following viral DNA entry which requires PIP_EF_ (23). Studies in *E. faecium* have identified cell wall polysaccharides, secreted antigen A, and RNA polymerase to be involved in phage infection (12, 24). Other studies have identified the enterococcal polysaccharide antigen (Epa) as a co-receptor for *E. faecalis* phages (25, 26).

Bacteria implement various mechanisms, including CRISPR-Cas and restriction-modification systems to resist phage infection (27). However, spontaneous mutation is the main mechanism driving both phage resistance and phage-bacteria coevolution (28). Phages must mutate to counter host mutations and persist in the population; their plastic genomes allow for the accumulation of adaptive mutations. (29, 30). Although there are numerous studies evaluating phage mutations during coevolution with laboratory strains of *Escherichia coli*, studies in medically relevant pathogens such as the enterococci are limited. A recent experiment coevolving *E. faecium* and myophage EfV-phi1 showed that phage tail fiber mutations helped overcome *E. faecium* phage resistance (12).

To further our understanding of phage-enterococcal interactions and their impact on genome evolution, we co-cultured the MDR *E. faecalis* strain SF28073 with two genetically distant phages, VPE25 and phage 47 (phi47) (23, 25), for 14 days with daily passaging. Both phages are long non-contractile tailed siphophages with double-stranded DNA genomes. Although both phages infect *E. faecalis* strain SF28073, nucleotide alignment revealed that their genomes only share 37.3% nucleotide identity, indicating they are genetically distinct. Orthologous protein clustering confirmed that these phages belong to unique enterococcal phage lineages. Based on these observations, we hypothesized that *E. faecalis* SF28073 may gain single nucleotide polymorphisms (SNPs) in cellular pathways and macromolecules that are specific to infection by either phage. To test this hypothesis, we ran two parallel co-culturing experiments. We show that *E. faecalis* SF28073 evolves unique mutations in response to either VPE25 or phi47 predation. We identified mutations in known macromolecules previously demonstrated to be necessary for *E. faecalis* phage infection; however, numerous undescribed mutations were also identified within a lower percentage of the *E. faecalis* population. Our work shows that surface associated factors are the major driver of *E. faecalis* phage resistance, yet genetic alterations emerge that implicate diverse metabolic pathways in the *E. faecalis* response to phage infection. Additionally, our data suggest that the ratio of phage to bacteria is an important factor when studying phage-bacterial co-evolution in vitro, as bottlenecks during serial passage may favor phage extinction.

## MATERIALS AND METHODS

### Routine bacterial culture

*E. faecalis* SF28073 (urine isolate from Michigan, USA) (31) was cultured in brain heart infusion (BHI, BD) medium at 37°C.

### Phage isolation and quantification

Bacteriophages VPE25 (23) and phage 47 (phi47) (25) were propagated using *E. faecalis* strains V583 (VPE25) or SF28073 (phi47) and phage titers were quantified by double agar overlay plaque assays, as described previously (23, 25). For clonal phage isolation, plaques were removed from agar overlays using a sterile p1000 pipette tip or a glass Pasteur pipette. Agar plugs were suspended in 1mL sterile SM-plus buffer (100mM NaCl, 50mM Tris-HCl, 8mM MgSO_4_, 5mM CaCl_2_ [pH 7.4]) and eluted overnight at 4°C. The eluted phages were filtered through a 0.45 μm syringe filter and stored at 4°C prior to phage titer determination by plaque assay.

### Coevolution assay

Individual colonies of *E. faecalis* SF28073 were grown overnight. The next day, 10^8^ colony forming units (CFU) of bacteria were inoculated into individual 125 mL flasks containing 25 mL of BHI broth supplemented with 10mM MgSO_4_. Five flasks were infected with 10^5^ plaque forming units (PFU) of phage VPE25 and five flasks with 10^5^ PFU of phi47, originating from individual plaques. Bacteria-only control cultures were included to identify mutations that arise due to laboratory passage in the absence of phage. All flasks were incubated at 37°C with shaking at 250 rpm. Every 24 hours, the cultures were passaged by transferring 250 μL of the culture to flasks containing 25 mL of fresh BHI media supplemented with 10mM MgSO_4_. At the time of passage, culture aliquots were removed for population DNA extraction and cryopreservation. For phi47, the culture media was centrifuged and filtered to isolate phages.

### DNA extraction for population sequencing

Genomic DNA was isolated from 1 mL culture aliquots using a previously described protocol for *E. faecalis* (32). Briefly, samples were treated with 5 mg/mL lysozyme for 30 minutes at 37°C. 0.5% SDS, 20mM EDTA and 50 μg/mL Proteinase K were added and incubated at 56°C for 1 hour. Samples were cooled to room temperature before adding an equal volume of phenol/chloroform/isoamyl alcohol and extracted by shaking. Samples were centrifuged at 17,000 rcf for 1 minute, and the aqueous layer was extracted with an equal volume of chloroform. Again, samples were centrifuged at 17,000 rcf for 1 minute, and nucleic acids were precipitated from the aqueous layer by adding 0.3M NaOAc [pH 7] and an equal volume of isopropanol. Nucleic acid was pelleted by centrifuging at 17,000 rcf for 30 minutes at 4°C, washed with 70% ethanol, and centrifuged at 17,000 rcf for 10 minutes. Finally, the pellet was dried and resuspended in sterile water. Genomic DNA was sequenced using Illumina the NextSeq 2000 platform to 300 Mbp depth at the Microbial Genome Sequencing Center (MiGS, Pittsburgh, PA, USA).

### Hybrid assembly of the *E. faecalis* SF28073 genome

The *E. faecalis* SF28703 genome was sequenced using Oxford Nanopore technology (ONT) as described previously (33, 34). Briefly, 1.5 μg genomic DNA was mechanically sheared into 8 kb fragments with a Covaris g-tube per the manufacturer’s instructions prior to library preparation with the ONT Ligation Sequencing Kit 1D (SQK-LSK108). Libraries were base called with MinKNOW (v3.5.5) to generate FASTQ and fast5 sequence reads. Illumina reads were obtained from MiGS as described above. Programs for DNA sequencing read processing and read assembly were run using the operating system Ubuntu 18.04.4 LTS. FASTQ sequences were filtered to gather reads with q scores >9 and length >1000 bp using Nanofilt (v2.5.0) (35). The adaptor sequences were trimmed from the filtered reads with Porechop (v0.2.3) (https://github.com/rrwick/Porechop). The processed MinION reads were co-assembled with Illumina reads using Unicycler (v0.4.7) with the default setting “normal mode” (34). Incomplete assemblies were manually completed as described in the “Unicycler tips for finishing genome” page (https://github.com/rrwick/Unicycler/wiki/Tips-for-finishing-genomes). Briefly, Bandage (v0.8.1) was used to visualize completion status of the assembly (36), and unassembled contig sequences were extracted. Using these unassembled contig sequences as baits, long reads from MinION sequences were gathered for incomplete regions using minimap2 (v2.11-r797) and an in-Bandage BLAST search was performed with the long reads against the graph (37). If long reads supported the continuity of two unassembled contigs, then the Bandage graph editing function was used to duplicate, delete edge, and merge contigs. The complete assembly sequence was saved from Bandage in FASTA format.

### Analysis of serially passaged bacterial populations using Illumina sequencing

Illumina reads from the bacterial populations obtained from MiGS were mapped to the assembled *E. faecalis* SF28073 chromosome (GenBank accession number CP060804) and the three endogenous plasmids (pSF1, CP060801; pSF2, CP060802; and pSF3, CP060803) using CLC Genomics Workbench with default settings. Detailed read mapping statistics were generated using the “QC for read mapping” tool in CLC Genomics Workbench with default settings to obtain the range of coverage and zero coverage regions in each assembly. The “Find low coverage” tool in CLC Genomics Workbench with the low coverage threshold set at 0 was used to manually inspect the regions found by the quality analysis to contain regions with 0 coverage. Sequence variants were identified using the “Basic variant detection” tool with a minimum coverage of 100, minimum frequency of 30%, and ploidy of 0. All variants identified were manually examined, and silent mutations were excluded from the analysis. Variants present in the bacteria-only controls were also excluded from further analysis.

### Phage 47 genome sequencing and analysis

Phi47 genomic DNA was isolated following the methods described above. A draft wild type phi47 genome was assembled *de novo* from Illumina reads obtained from MiGS, with the largest contig forming the full genome. RAST genome annotation (v2.0) was used to predict gene function (38). During coevolution with *E. faecalis* SF28073, culture media was filtered through a 0.45-μm filter. DNA was extracted from filtered media using the proteinase K and phenol/chloroform method described above and sequenced using Illumina technology at MiGS. These reads were mapped to the wild type phi47 genome and SNPs were identified using CLC Genomic Workbench with a minimum coverage of 10, a minimum frequency of 30%, and ploidy of 0.

### OrthoMCL analysis

Enterococcal phage phylogeny was determined using OrthoMCL (39) as described previously (24). Enterococcal phage genomes were downloaded from the INPHARED phage genome database (18). As of July 1, 2021, there were 126 enterococcal phage genome sequences available in addition to our inclusion of the phi47 genome. Proteomes determined using Prodigal (40) were used as input into an OrthoMCL MySQL database. A cluster inflation value of 1.5 was used and the resulting matrix was input for ggdendro and ggplot2 packages in R version 3.6.3. The dendrogram was determined using the average linkage method for hierarchical clustering of Manhattan distance metrics.

### Data Availability

The *E. faecalis* SF28073 chromosome and its three endogenous plasmids can be in the NCBI database under the following accession numbers; *E. faecalis* SF28073 chromosome (CP060804), *E. faecalis* SF28073 plasmids (pSF1, CP060801; pSF2, CP060802; and pSF3, CP060803). Illumina DNA sequencing reads associated with this study are deposited at the European Nucleotide Archive under accession number PRJEB48380.

## RESULTS

### Phi47 and VPE25 phages are genetically distinct

phi47 depends on the enterococcal polysaccharide antigen (Epa) for adsorption to host cells (25). The phi47 genome is 57,289 base pairs in length, consisting of 101 predicted open reading frames (ORFs). Using RAST genome annotation (38), we characterized the phi47 genome based on functional classifications (Fig. 1A). The genome exhibits typical modularity; meaning that tail, structural, and DNA replication genes are in proximity to genes of similar function. The remainder, and the majority of the genes, are predicted to be hypothetical.

**Figure 1:**
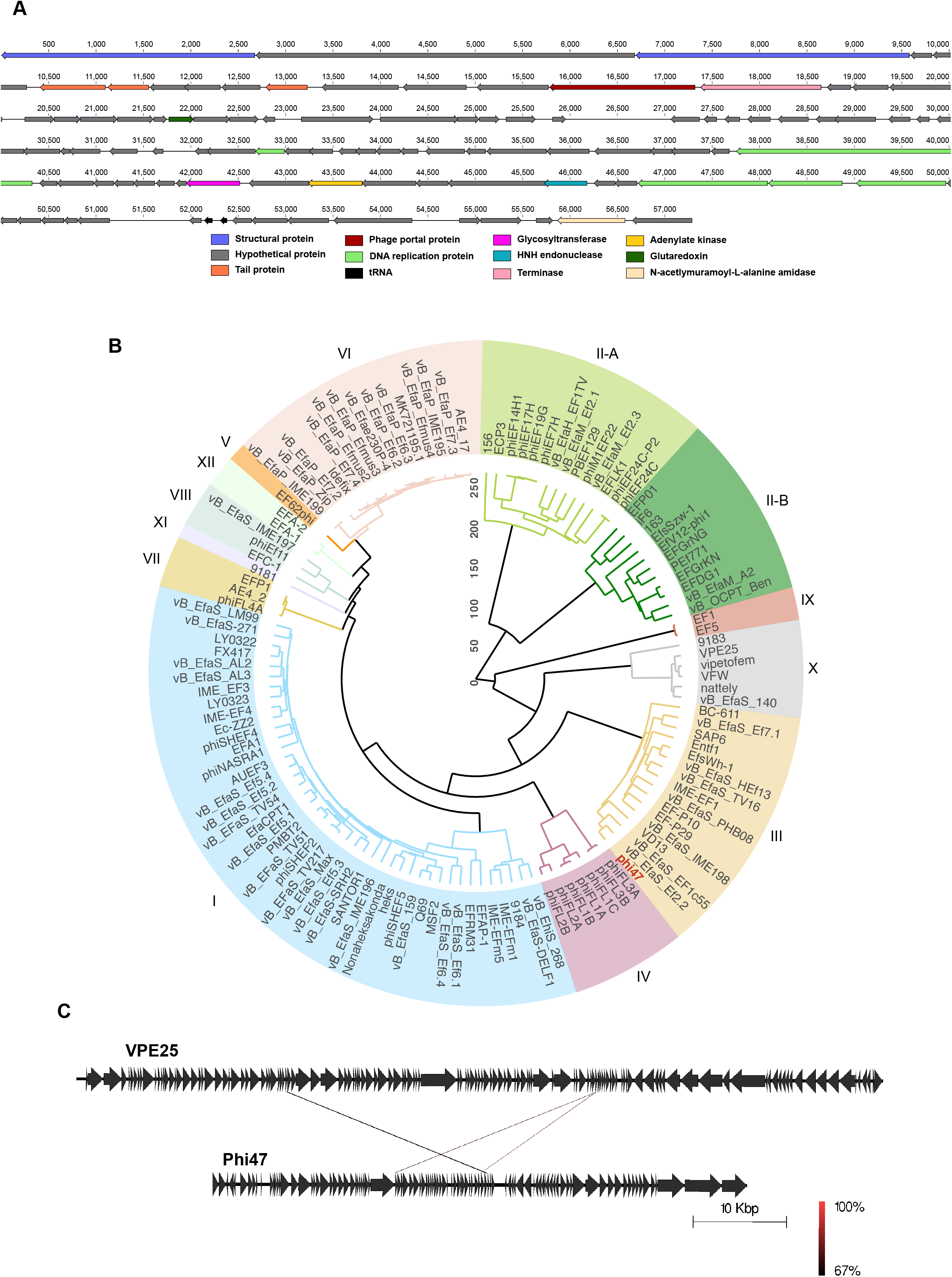
Phi47 and phage VPE25 are genetically distinct. **(A)** RAST genome annotation predicted the function of 16 open reading frames out of 101, and 2 tRNAs. Open reading frames encoding proteins of similar function are depicted in the same color. **(B)** Comparative analysis shows phage 47 belongs to orthocluster III. OrthoMCL was used to compare the phi47 genome to all publicly available enterococcal phage genomes. A phylogenetic proteomic tree was generated from OrthoMCL. Height is the average linkage of hierarchical clustering with 1000 iterations using the Manhattan distance metric. 126 enterococcal phage genomes from the INPHARED database were used for comparison to phi47 (in red text). Distinct phage orthoclusters are represented by colored boxes. Roman numerals next to shaded boxes designate the orthocluster number. **(C)** EasyFig analysis shows three genes shared between phage 47 and VPE25. These genes are 67-100% identical at the nucleotide level.

Phi47 and VPE25 are both siphoviruses belonging to the class *Caudoviricetes* (23, 25, 41). However, these phages differ in host range and genome content. VPE25 is a virulent phage capable of infecting numerous *E. faecalis* strains (23), including SF28073, while phi47 primarily infects SF28073 (25). Comparative genomic analysis of phi47 was performed with all publicly available enterococcal phage genomes using OrthoMCL (Fig. 1B) (18). This algorithm generates a phylogenetic tree of clustered phage genomes (orthoclusters) based on orthologous proteins (24, 39, 42). Of the 12 known orthoclusters (24, 42), phi47 belongs to cluster III, while VPE25 is in cluster X. Further analysis of orthocluster II resulted in its division into two unique orthoclusters bringing the total to 13 orthoclusters of enterococcal phages (Fig 1B). EasyFig comparison of the phage genomes revealed only three shared genes (gray lines, Fig. 1C) (43). These genes exhibit 67% or greater identity at the nucleotide level. Together, these genetic analyses demonstrate the lack of common genes between phages VPE25 and phi47, making them genetically distinct.

### Phage infection of *E. faecalis* promotes mutations in cell wall macromolecules necessary for phage infection, and unique mutations accumulate in a phage-dependent manner

To identify bacterial mutations that confer phage resistance in *E. faecalis* SF28073, we conducted two independent coevolution experiments, infecting five replicate SF28073 cultures with phages derived from individual plaques of either phage VPE25 or phi47, and passaged these cultures for 14 consecutive days. Bacteria-only controls were established and treated under identical conditions in the absence of phage infection. Genomic DNA from the bacterial populations were sequenced from each replicate at five time points (days 0, 1, 3, 7 and 14). To identify mutations in the SF28073 genome, sequencing reads were mapped to the closed *E. faecalis* SF28073 reference genome generated in this study by hybrid assembly of Illumina and Oxford Nanopore MinION sequencing reads. The assembled SF28073 genome consists of the chromosome and three endogenous plasmids designated pSF1, pSF2, and pSF3 (GenBank accession numbers CP060804, CP060801, CP060802, and CP060803, respectively).

Non-synonymous, unique bacterial single nucleotide polymorphisms (SNPs) were observed in all experimental replicates, except in one of the VPE25-challenged replicates where the mutation frequencies were below our 30% population-wide cutoff. Interestingly, the mutations that arose in *E. faecalis* SF28073 challenged with phage VPE25 largely differed from mutations in *E. faecalis* SF28073 challenged with phi47. As expected, we observed mutations in PIP_EF_ at one or more time points in all of the VPE25 challenged replicates, except the culture mentioned above which did not meet our read mapping cutoff (Table S1). We detected *epa* mutations in 4 of the cultures infected with phi47 (Table S2). These two macromolecules have been previously reported to be essential for successful infection of these phages; the integral membrane protein PIP_EF_ is the receptor for VPE25, while both VPE25 and phi47 rely on the enterococcal polysaccharide antigen (Epa) for adsorption (23, 25, 44).

We identified *ccpA* as the only common gene mutated when SF28073 was challenged with either VPE25 or phi47 (Table 1, 2). When exposed to phage VPE25, *ccpA* had mutations in two replicates which appeared at different time points (days 7 and 14, Table S1), while exposure to phi47 resulted in one replicate harboring a *ccpA* mutation on day 14 (Table S2). In *E. faecalis*, catabolite control protein A (CcpA) plays a key role in regulating transcription of proteins involved in carbon source utilization (45). Moreover, both experimental groups had mutations arise in different components of the SUF system, which is involved in the iron-sulfur (Fe-S) cluster assembly pathway (46) (Table S1, S2). One VPE25-challegend replicate had a mutation in *sufU*, encoding a sulfur relay protein (46), while one phi47-challenged replicate had a mutation in *sufD*. Both mutations were identified on day 7 and were maintained to day 14. This suggests that both phages may utilize bacterial iron-sulfur complexes during infection.

A mutation in the mannose-permease encoding gene *manX* was found in one replicate infected with phage VPE25. In a previous study, mutations in *E. coli* subunits of the ManXYZ mannose-permease were mutated after infection with phage Lambda. This component of the phosphotransferase system is known to be used by phage Lambda to eject its DNA (29), suggesting that phage VPE25 may implement a similar infection mechanism by interacting with the mannose phosphotransferase system to infect *E. faecalis* SF28073.

Additionally, one replicate challenged with phage VPE25 had mutations in a putative restriction-modification (R-M) system, located within the two specificity (S) subunit genes, specifically genes H9Q64_13860 and H9Q64_13845 (Table S1). In R-M systems, the S subunit genes, composed of two target recognition domains, recognize specific DNA sequences thereby providing target specificity to the R-M complex (47). On day 7, both subunits shared missense mutations resulting in amino acid changes from leucine to phenylalanine and lysine to glutamine. Surprisingly, on day 14 these mutations were no longer detected, suggesting that they rendered these cells less fit in the population. Other missense mutations specific to each S subunit gene were also found on day 7. These two mutations were observed again on day 14 at higher frequencies, which on the contrary, suggest that these mutations provided a fitness advantage to the population. Lastly, day 14 revealed two additional mutations in both S subunits that were not present on day 7. The numerous amino acid changes observed across the S subunits of the R-M system suggests these mutations may be increasing the specificity of the subunit S towards recognizing the VPE25 genome.

When *E. faecalis* SF28073 coevolved with phi47, we observed three genes mutated across multiple replicates (Table 2). H9Q64_01755, a predicted transposase, was mutated in two replicates. In fact, both replicates had the same mutation arginine 144 to leucine. H9Q64_09795 was also mutated in two replicates. This gene, *epaAC*, is a predicted epimerase/dehydratase (48). Lastly, H9Q64_09850 was mutated in three replicates. This gene is *epaR*, the final gene in the rhamnose-sugar biosynthesis locus of *epa* that is a predicted priming glycosyltransferase (48). Epa is involved in phage adsorption (25, 26), making this gene essential for successful phi47 infection.

### Phi47 co-evolves mutations in tail and hypothetical genes

During the 14 days of passaging, we enumerated both phi47 and *E. faecalis* SF28073 to determine the population kinetics for each experimental replicate (Fig. 2). We observed different phage abundance patterns across the five replicates, despite each being treated identically. While all replicates had an expected spike in phi47 titer on day 1 after 24 hours of replication in a completely susceptible population, and a reduction in titer on day 3, phage abundance differed for each replicate on day 7 (Fig. 2). Culture 1 phi47 titer spiked and was followed by a continuous decline until phi47 was no longer detectable in the culture via plaque assay by day 11. Cultures 2 and 3 had no detectable phi47 on day 7 and became extinct. Cultures 4 and 5 had low phage titers on day 7. Culture 4 had a phi47 spike on day 9 followed by a decline until it was no longer detected on day 12, and phi47 went extinct in culture 5 by day 9.

**Figure 2:**
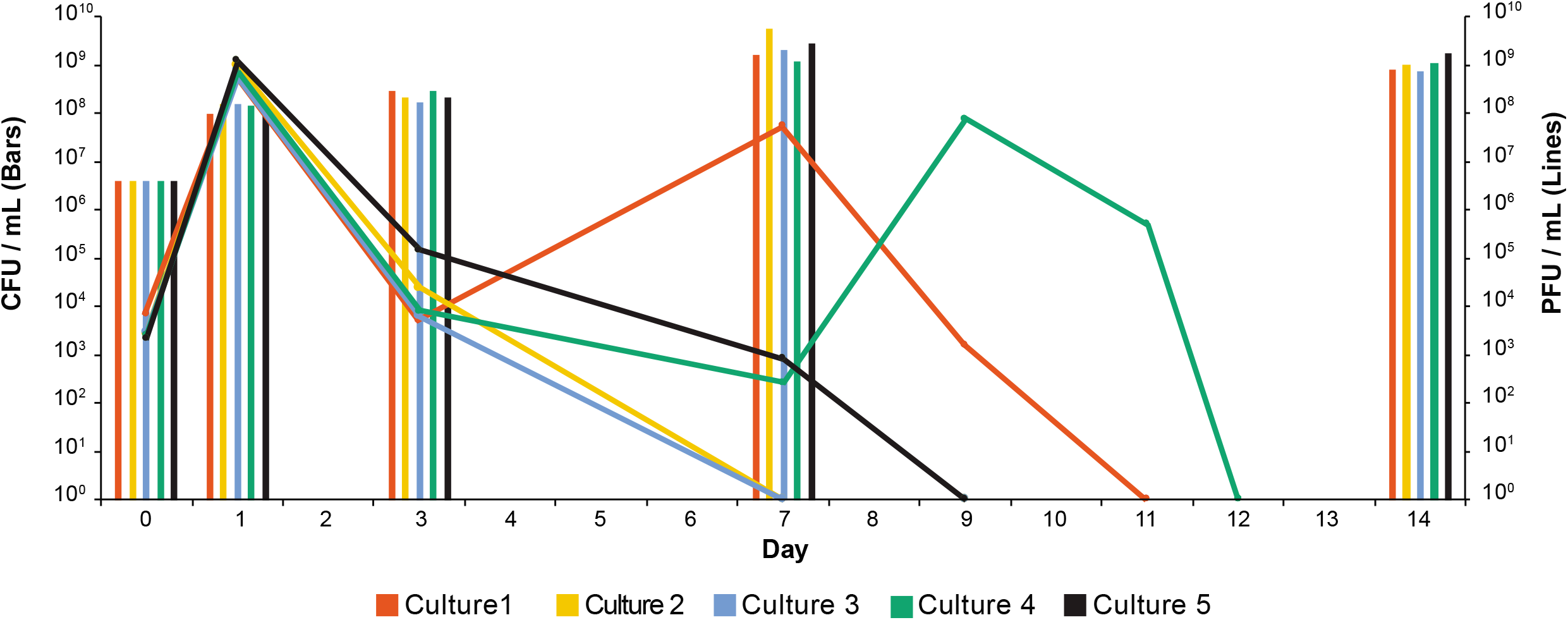
Phi47 kinetics differ in each experimental replicate. Lines and bars of the same color represent *E. faecalis* SF28073 and phi47 titers, respectively. Bars represent the mean of 3 technical replicates, while lines are a single biological replicate. Additional plaque assays were performed to identify when phi47 became undetectable in each culture.

Due to the differences observed in phi47 abundance over the course of these experiments, we sequenced the phage population of each replicate on days 0, 1, 3, and 7. Day 14 was excluded from analysis because no replicate had phage detectable by plaque assay at that time point. Sequencing of phage DNA revealed that each culture had a unique mutation profile. Phages from cultures 2 and 3 acquired the most SNPs, but viable phages were undetectable at day 7, suggesting that these acquired mutations in various tail and hypothetical proteins may have been deleterious to the phages’ ability to overcome bacterial resistance mutations. Phages from cultures 3 and 4 developed identical tail SNPs on day 7. While there were no infectious phages detectable by plaque assay in culture 3 on day 7, we were still able to recover phage DNA in culture media, allowing us to perform genetic analyses. Culture 5 phages only developed one SNP in a minor structural protein on day 3, which was maintained on day 7. Culture 1 phages developed no SNPs. Interestingly, there are three hypothetical genes that are mutated in phages across multiple cultures (Table S3). Despite the different mutations observed across replicates, phi47 was not detectable in any of the cultures by the end of the passaging, indicating that these phages were unable to subvert phage resistance leading to their extinction.

## DISCUSSION

*E. faecalis* is a commensal and nosocomial pathogen, and is becoming increasingly resistant to last resort antibiotics (49). In this study, we show the coevolution of *E. faecalis* SF28073 with two genetically distinct phages, VPE25 and phi47. Our results reveal that the *E. faecalis* genome mutates in district patterns in response to each phage. This indicates that genetically unique phages elicit distinct genetic responses within the same host. In particular, for both phages, *E. faecalis* developed missense mutations in cell wall macromolecules, specifically PIP_EF_ and Epa, that are required for successful infection by VPE25 and phi47, respectively (23, 25). VPE25 has recently been shown to depend on Epa (44), most likely for adsorption to the cells. Because PIP_EF_ and Epa are essential for successful phage infection, mutations in these genes prevent phage infection. Despite this, we observed no *epa* mutations in cultures challenged with VPE25, suggesting that PIP_EF_ mutations are dominant and potentially more advantageous than *epa* mutations, likely due to the fitness costs caused by *epa* mutations (25, 26). SNPs in *pip_EF_* and *epa* indicate that across bacterial species, phage receptor and co-receptor mutations are common to prevent phage infection.

Mutations in *ccpA* were found in cultures challenged with both VPE25 and phi47. In Gram-positive bacteria, CcpA regulates the expression of genes encoding proteins involved in the catabolism of complex carbon sources when more rapidly metabolized carbohydrates such as glucose or fructose are present (45, 50). Phage infection outcome is determined by the bacterial host physiological condition (51). For instance, type of carbon source present in the bacterial host’s media affects phage development and their ability to lyse the cells (51, 52). It is possible that mutations in *ccpA* could impact bacterial carbon source utilization, allowing the bacteria to switch to using carbohydrates less preferred by the phages, therefore negatively impacting phage production. Furthermore, Chatterjee et al. showed that RNA sequencing of OG1RF_10887 was over-expressed early during VPE25 infection (44). Here, we observed mutations in the SF28073 iron-sulfur binding genes *sufD* and *sufU* during phi47 and VPE25 infection, respectively. These two studies suggest that iron-sulfur binding proteins are critical during lytic phage infection.

Recent work in *E. coli* has revealed mutations in OmpF (a phage Lambda receptor) and in subunits of the *manXYZ* mannose permease operon, both of which phage Lambda uses for DNA ejection into the host cytoplasm. We discovered a mutation in the mannose permease gene *manX*, a component of the mannose phosphotransferase system, in one of our cultures infected with phage VPE25, suggesting that phage VPE25 could also utilize a similar infection process by interacting with the mannose phosphotransferase system to infect *E. faecalis*.

Additionally, we show that phi47, an understudied enterococcal phage, develops mutations during co-culture with its host. Phages from cultures 3 and 4 developed identical mutations in the major tail protein on day 7. However, culture 3 had no phages detectable by plaque assay on that day, suggesting that these phages were unable to overcome the bacterial mutations in cell wall associated macromolecules, such as Epa. Phages in culture 4 developed the same major tail protein mutations before the bacteria developed *epa* locus mutations, suggesting that the major tail protein mutations arose independent of host *epa* mutations. Despite these novel mutations, phi47 could not be maintained in the experimental cultures.

While there is currently only one paper investigating phage and *Enterococcus* coevolution (12), there are numerous coevolution studies between the model organism *E coli* and its phages. In one study *E. coli* and phage T3 were coevolved in chemostats, allowing for a controlled experimental environment. Under this condition, the authors observed common bacterial mutations at the gene level and phage mutations at the codon level across experimental replicates (53). We believe that our current study both supports and contradicts these findings. In our study, four of the five cultures challenged with phage VPE25 developed mutations in PIP_EF_. This maintains the conclusion that bacterial mutations in response to phage pressure reproducibly occur at the gene level. However, cultures challenged with phi47 showed more variability among *E. faecalis* genomic mutations across replicates. While phi47 developed some mutations that were shared across multiple cultures, each culture had a unique phage SNP profile, which are at odds with Perry et al. that phage mutations replicate at the codon level. It is possible that these outcomes may depend on the phage-host bacterial pair used in coevolution experiments, the specific MOI used to initially infect the cultures, and the growth conditions tested.

Ultimately, the study by Perry et al. suggests that lack of reproducibility in coevolution experiments may be due to abiotic selection pressures, meaning that divergence among experimental replicates can increase via random events occurring in each population over time. This argues for a more controlled and consistent experimental design, such as a chemostat for continuous culturing, that may reduce stochastic events. We believe that our experimental design, which included manual daily sub-culturing, may have introduced bottlenecks causing a selection bias for the growth and preservation of bacteria, while causing phages to disappear from the population. In our experiments, we initially inoculated each flask with ~10^8^ CFU of *E. faecalis* SF28073 and ~10^5^ PFU of phage (MOI 0.001). A similar ratio (MOI 0.003) that was used to study phage-*E. faecium* coevolution (53). However, for the *E. faecium* study, passaging was performed at a 1:10 ratio, transferring every 12 hours for a total of 16 time, while we implemented a passage ratio of 1:100, transferring every 24 hours for 14 days.

A study in *E. coil* used a chemostat with an MOI of 2, showing that in this setting coevolution happened in the form of adaptation and counter-adaptation (54). While we began to observe phage extinction by day 7, the phage population from Wandro et al. was maintained in all cultures throughout the course of the experiment. We speculate that the small volume we sub-cultured, the low starting MOI, and the fact that there were magnitudes more bacterial cells than phage in the population, may have caused a significant decrease in the number of phages passaged, thus introducing a bottleneck that ultimately eliminated the phage from the population. Additionally, our 24-hour passages may have allowed extra time for bacteria to grow and continue mutating to resist phage infection.

Our study highlights the importance of carefully considering experimental design factors such as MOI and sub culturing methods when studying the coevolution of phages and their hosts to prevent the introduction of bottlenecks. Studies such as transposon-insertion sequencing used to investigate genes involved in infection can be confounded by the presence of bottlenecks that can limit the quality of the library (55). Furthermore, bottlenecks could prevent the discovery of novel genes involved in phage infection by limiting the survival of the phage in the population. Future studies should consider our methods and modify them to support continuous phage replication—for instance, using higher volumes if manually passaging, implementing a higher starting MOI, and the use of continuous culturing systems.

## ACKNOWLEDGEMENTS

This work was supported by National Institutes of Health grants R01AI141479 (B.A.D.), R01AI116610 (K.L.P.), T32AI052066 (C.N.J.), F31AI157050 (C.N.J.), and T32AR007534 (M.R.M.).

---

**Supplemental table 1:** All mutations present in *E. faecalis* SF28073 challenged with VPE25.

**Supplemental table 2:** All mutations present in *E. faecalis* SF28073 challenged with phi47.

**Supplemental table 3:** All mutations present in phi47 coevolved with *E. faecalis* SF28073.

