## Supplemental Table 1 for "Genetically distant bacteriophages elicit unique genomic changes in *Enterococcus faecalis*"

| Culture | Gene | Predicted function | AA Change | Frequency |
|---|---|---|---|---|
| **Day 1** | | | | |
| 1,2,4 | H9Q64_06285 | Phage Infection Protein (PIP) | Ser36* | 99.4%, 96.0%, 86.7% |
| **Day 3** | | | | |
| 2 | H9Q64_06285 | Phage Infection Protein (PIP) | Gln462* | 33.0% |
| | H9Q64_15390 | PTS mannose/fructose/sorbose transporter family subunit IID | Asn88fs | 33.3% |
| **Day 7** | | | | |
| 1 | *manx* | ManX | Asn77Lys | 48.8% |
| | H9Q64_06285 | Phage Infection Protein (PIP) | Ser36* | 98.6% |
| 2 | H9Q64_13860 | Restriction endonuclease subunit S | Leu166Phe, Lys180Glu, Thr160Ala | 30.0%, 38.7%, 40.0% |
| | H9Q64_13845 | Restriction endonuclease subunit S | Leu181Phe, Lys180Glu, Ala175Thr | 32.3%, 36.5%, 38.9% |
| 4 | H9Q64_06285 | Phage Infection Protein (PIP) | Ser685fs | 55.9% |
| | *ccpA* | Catabolite control protein A | Asn27Ile | 57.5% |
| 5 | H9Q64_06285 | Phage Infection Protein (PIP) | Ile739fs | 39.2% |
| | *sufU* | SufU, component of the SUF system | Thr70Ile | 39.5% |
| **Day 14** | | | | |
| 1 | *ccpA* | Catabolite control protein A | Tyr94Asn | 32.5% |
| | *manx* | ManX | Asn77Lys | 42.5% |
| | H9Q64_06285 | Phage Infection Protein (PIP) | Ser36* | 95.5% |
| 2 | H9Q64_13860 | Restriction endonuclease subunit S | Asn157Asp, Thr160Ala | 46.5%, 56.4% |
| | H9Q64_13845 | Restriction endonuclease subunit S | Asp172Asn, Ala175Thr | 42.7%, 82.6% |
| 5 | H9Q64_06285 | Phage Infection Protein (PIP) | Ile739fs | 71.3% |
| | *sufU* | SufU, component of the SUF system | Thr70Ile | 71.7% |
