## Supplemental Table 2 for "Genetically distant bacteriophages elicit unique genomic changes in *Enterococcus faecalis*"

| Culture | Gene | Predicted function | AA Change | Frequency |
|---|---|---|---|---|
| **Day 1** | | | | |
| None detected | | | | |
| **Day 3** | | | | |
| 4 | H9Q64_01755 | Transposase | Arg144Leu | 97.0% |
| **Day 7** | | | | |
| 1 | H9Q64_09795 | Epimerase, *epaAC* | Glu208* | 66.0% |
| | *sufD* | FeS assembly protein | Arg295Gln | 66.5% |
| | *ptsP* | Phosphoenolpyruvate-protein phosphotransferase | Asp267Ala | 70.0% |
| 2 | H9Q64_09795 | Epimerase, *epaAC* | Gly191Asp | 31.0% |
| 4,5 | H9Q64_01755 | Transposase | Arg144Leu | 33.5%, 34.5% |
| **Day 14** | | | | |
| 1 | *ccpA* | Catabolite control protein A | Leu306Phe | 33.5% |
| | *ptsP* | Phosphoenolpyruvate-protein phosphotransferase | Asp267Ala | 31.6% |
| | H9Q64_08040 | DUF1189 domain-containing protein | Ala133Thr | 41.8% |
| | H9Q64_09795 | Epimerase, *epaAC* | Glu208* | 36.0% |
| | H9Q64_09850 | Sugar transferase, *epaR* | Pro320Leu | 43.3% |
| | *sufD* | FeS assembly protein | Arg295Gln | 31.4% |
| 2 | H9Q64_09795 | Epimerase, *epaAC* | Gly191Asp | 43.3% |
| 3 | H9Q64_09850 | Sugar transferase, *epaR* | Thr296Ile | 43.4% |
| 5 | H9Q64_06955 | Directly upstream of *ptsP*, phosphocarrier protein HPr | Ala16Val | 44.4% |
| | H9Q64_09850 | Sugar transferase, *epaR* | Pro320Leu | 52.0% |
