## Supplemental Table 3 for "Genetically distant bacteriophages elicit unique genomic changes in *Enterococcus faecalis*"

| Culture | Position | Reference | Allele | Annotation | AA Change | Frequency |
|---|---|---|---|---|---|---|
| **Day 1** | | | | | | |
| None detected | | | | | | |
| **Day 3** | | | | | | |
| 2 | 7802 | - | T | Minor capsid protein | Thr594fs | 100% |
| | 21559 | A | T | Hypothetical protein | Glu112Val | 100% |
| | 21562 | G | T | | Arg113Leu | 100% |
| | 29876 | G | A | Hypothetical protein | Pro122Leu | 100% |
| | 31004 | C | - | Hypothetical protein* | Val14fs | 38.8% |
| | 31009 | - | A | | Lys12fs | 55.5% |
| | 31285 | C | G | Hypothetical protein | Arg51Thr | 98.9% |
| | 34228 | C | - | Hypothetical protein* | Gly59fs | 52.1% |
| | 35922 | A | - | Hypothetical protein | Ser94fs | 100% |
| | 41040 | A | - | Hypothetical protein | Tyr49fs | 100% |
| | 41444 | T | - | Hypothetical protein | Asn59fs | 91.3% |
| | 44217 | - | A | Hypothetical protein | Gln54fs | 93.3% |
| | 45366 | C | G | Hypothetical protein* | Glu126Asp | 100% |
| | 45368 | C | G | | Glu126Gln | 100% |
| | 45373 | A | T | | Met124Lys | 100% |
| | 55247 | - | C | Hypothetical protein | Arg63fs | 94.4% |
| 3 | 31004 | C | - | Hypothetical protein* | Val14fs | 63.1% |
| | 31009 | - | A | | Lys12fs | 36.8% |
| | 31040 | T | - | | Met2fs | 50.0% |
| | 48552 | A | - | DNA replication protein | Asn108fs | 68.7% |
| | 48556 | - | G | | Lys106fs | 68.7% |
| 4 | 17578 | - | C | Phage terminase, large subunit | Tyr358fs | 22.2% |
| 5 | 100 | A | C | phage minor structural protein | Val859Gly | 94.9% |
| **Day 7** | | | | | | |
| 2 | 27651 | - | TG | Hypothetical protein | Asp48fs | 66.6% |
| | 27652 | - | T | | Asp48fs | 66.6% |
| | 27655 | T | C | | Asn46Ser | 66.6% |
| | 27657 | C | A | | Met45Ile | 66.6% |
| | 31004 | C | - | Hypothetical protein* | Val14fs | 43.7% |
| | 31009 | - | A | | Lys12fs | 52.9% |
| | 31040 | T | - | | Met2fs | 53.3% |
| | 34228 | C | - | Hypothetical protein* | Gly59fs | 42.1% |
| 3 | 10451 | C | T | Major tail protein | Val217Ile | 100% |
| | 10453 | G | A | | Ala216Val | 100% |
| | 10457 | - | C | | Thr215fs | 100% |
| | 10459 | T | C | | Asn214Ser | 100% |
| | 10463 | C | - | | Val213fs | 100% |
| | 28104 | C | T | Hypothetical protein | Glu4Lys | 100% |
| | 28107 | G | A | | Pro3Ser | 100% |
| | 31040 | T | - | Hypothetical protein* | Met2fs | 55.5% |
| | 34228 | C | - | Hypothetical protein* | Gly59fs | 34.7% |
| | 45366 | C | G | Hypothetical protein* | Glu126Asp | 100% |
| | 45368 | C | G | | Glu126Gln | 100% |
| | 45373 | A | T | | Met124Lys | 100% |

| | | | | | | |
|---|---|---|---|---|---|---|
| 4 | 10451 | C | T | Major tail protein | Val217Ile | 37.5% |
| | 10453 | G | A | | Ala216Val | 37.5% |
| | 10457 | - | C | | Thr215fs | 35.2% |
| | 10459 | T | C | | Asn214Ser | 35.2% |
| | 10463 | C | - | | Val213fs | 33.3% |
| | 17578 | - | C | Phage terminase, large subunit | Tyr358fs | 50.0% |
| | 20510 | G | A | Hypothetical protein | Arg88Lys | 27.7% |
| 5 | 100 | A | C | Phage minor structural protein | Val859Gly | 93.6% |

* Indicates genes mutated across multiple cultures.
